# Application of a gut-immune co-culture system for the study of N-glycan-dependent host-pathogen interactions of *Campylobacter jejuni*

**DOI:** 10.1101/702472

**Authors:** Cristina Y. Zamora, Elizabeth M. Ward, Jemila C. Kester, Wen Li Kelly Chen, Jason G. Velazquez, Linda G. Griffith, Barbara Imperiali

## Abstract

An in vitro gut-immune co-culture model with apical and basal accessibility, designed to more closely resemble a human intestinal microenvironment, was employed to study the role of the Nlinked protein glycosylation (Pgl) pathway in Campylobacter jejuni pathogenicity. The gutimmune co-culture (GIC) was developed to model important aspects of the human small intestine by the inclusion of mucin producing goblet cells, human enterocytes, and dendritic cells, bringing together a mucus-containing epithelial monolayer with elements of the innate immune system. The utility of the system was demonstrated by characterizing host-pathogen interactions facilitated by N-linked glycosylation, such as host epithelial barrier functions, bacterial invasion and immunogenicity. Changes in human intestinal barrier functions in the presence of 11168 C. jejuni (wildtype) strains were quantified using GICs. The glycosylationdeficient strain 11168 ΔpglE was 100-fold less capable of adhering to and invading this intestinal model in cell infectivity assays. Quantification of inflammatory signaling revealed that 11168ΔpglE differentially modulated inflammatory responses in different intestinal microenvironments, suppressive in some but activating in others. Virulence-associated outer membrane vesicles produced by wildtype and 11168ΔpglE C. jejuni were shown to have differential composition and function, with both leading to immune system activation when provided to the gut-immune co-culture model. This analysis of aspects of C. jejuni infectivity in the presence and absence of its N-linked glycome, is enabled by application of the gut-immune model and we anticipate that this system will be applicable to further studies of C. jejuni and other enteropathogens of interest.

## Introduction

The Gram-negative pathogen *Campylobacter jejuni* is a leading cause of gastroenteritis and diarrheal disease. Infections can be severe, especially in children, the elderly and immunocompromised individuals, with infection leading to fatality in 1 in 3,000 cases by some estimates (Ternhag et al. 2005). It has been suggested that adhesion to and invasion of host epithelial cells is critical for disease development (Young, 2007). *C. jejuni* has been shown to transcellularly invade intestinal epithelial cells, becoming encased in vacuoles that protect them from lysosomal degradation and immune detection (Dorrell et al. 2001) (Watson and Galan 2008). Characterization of the roles of virulence determinants in adhesion and invasion is crucial for identification of pathways, enzymes, and molecules to target for therapeutic intervention. This is of particular importance in the era of antimicrobial resistance, where additional insight into the contributors to microbial pathogenicity would allow for investigation of anti-virulence agents that may be less likely to elicit such resistance. Many virulence factors have been associated with host cell invasion, including flagellar motility and chemotaxis, outer membrane vesicles, adhesins and proteases.

One such determinant of microbial pathogenicity is protein glycosylation (Lu et al. 2015). The global significance of N-linked glycosylation in *C. jejuni* is not yet fully understood, however a correlation between N-linked protein glycosylation and virulence has been demonstrated in avian models of infection and *in vitro* mammalian systems. The N-linked protein glycosylation (*pgl*) operon and N-glycan of *C. jejuni* are illustrated in Fig 1A. Cecal analysis of chick intestines has shown significantly-decreased *C. jejuni* colonization upon individual knockouts of *pglB, pglD, pglE, pglH* and *pglK* genes (Jones et al. 2004, Kelly et al. 2006, Szymanski et al. 2002). Additionally studies of 81-176 *C. jejuni* in murine models have revealed similar decreases in colonization by *pglB* and *pglE* knockouts (SzymanskiBurr et al. 2002). *In vitro* studies of the Pgl pathway and its role in virulence have also been carried out in unpolarized monolayers of HeLa (MacCallum et al. 2005), INT 407 (MacCallumHaddock et al. 2005), T84 MacCallumHaddock et al. (2005), and mucus-secreting HT29 (Bahrami et al. 2011, Guerry et al. 2002). However, these systems do not feature tight junctions, brush borders or other important components of intestinal epithelia (Backert et al. 2013). Additionally, studies of infectivity on polarized monolayers of human epithelial colorectal adenocarcinoma (Caco-2) (Hu et al. 2008) do not account for the intestinal mucosal layer and its barrier functions, which in some strains of *C. jejuni* has been shown to play a role in bacterial motility (Ferrero and Lee 1988) and assist in binding and invasion of host cells (Szymanski et al. 1995). Bacterial interactions with mucosal components of the gut have also been shown to be an important part of virulence behaviors, in some cases influencing whether *C. jejuni* acts as a commensal as seen in chickens and other animals, or as a pathogen as seen in humans(Alemka et al. 2010, Alemka et al. 2012). Although these *in vitro* studies on components of human intestinal physiology have provided valuable insight, a more comprehensive understanding of the mechanism and determinants of *C. jejuni* pathogenicity could be potentially achieved with model systems incorporating several aspects and more closely resembling human gut physiology.

**Fig 1.**
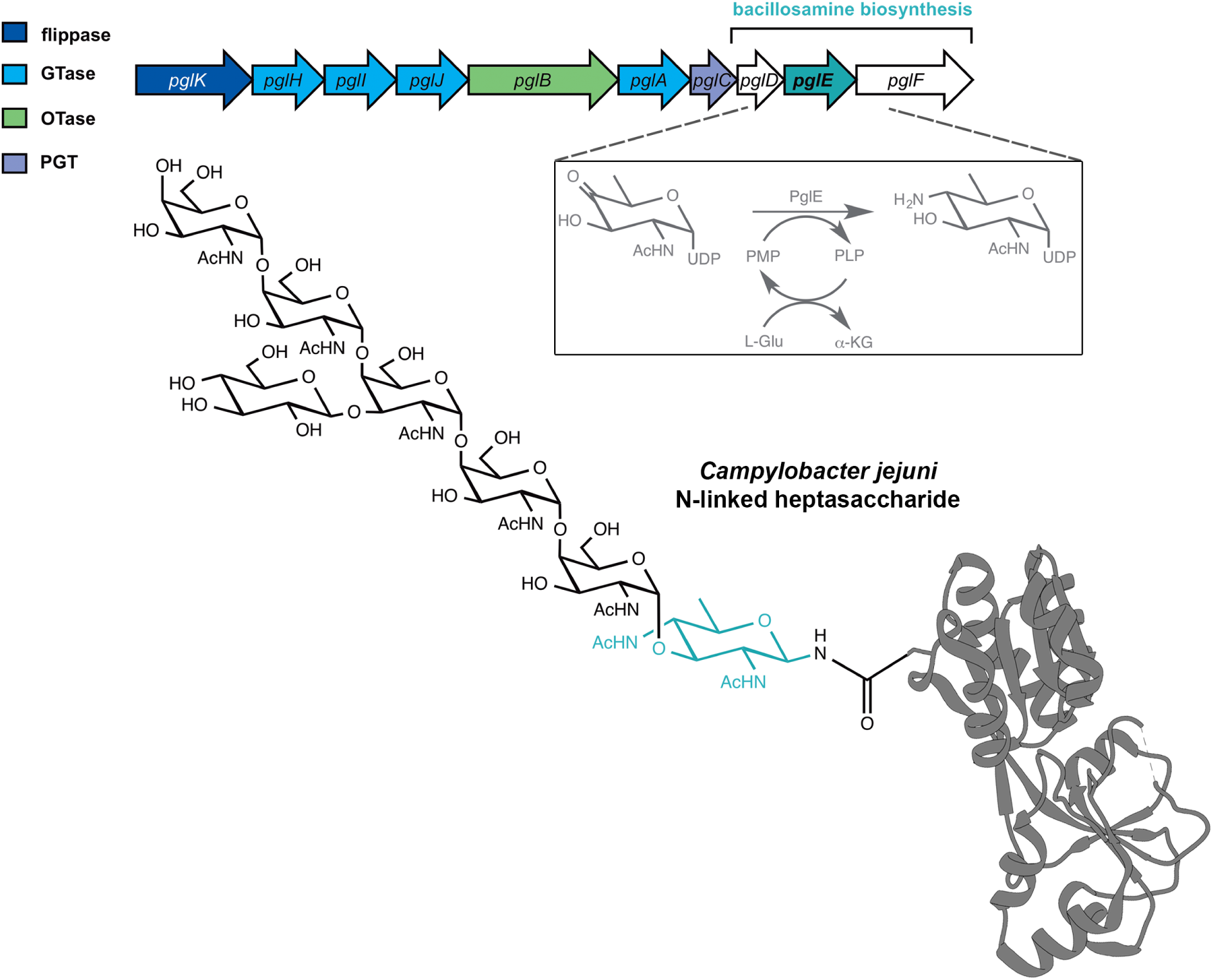
Illustration of the pgl locus in *C. jejuni* and the resulting N-linked heptasaccharide. Ten enzymes act in concerted fashion to produce N-linked glycoproteins in *C. jejuni*. Knockout of transaminase PglE results in near-complete loss of function in this biosynthetic pathway. Shown as a cartoon is putative adhesin PEB3 from *C. jejuni* (PDB: 2HXW), N-glycosylated with the resulting heptasaccharide.

To address this need, herein we present the adaptation and application of a gut-immune co-culture model (GIC, Figure 2), engineered to mirror important aspects of native human small intestinal tissue including barrier functions and pharmacological properties (Chen et al. 2017, Edington et al. 2018, Louwen et al. 2012, Tsamandouras et al. 2017), for the characterization of *C. jejuni* N-linked glycans in host-pathogen interactions. Each GIC contains an epithelial monolayer comprising a 9:1 ratio of absorptive human enterocytes (C2BBe1) and human mucin-producing goblet cells (HT29-MTX) seeded and grown on a permeable Transwell insert. The C2BBe1 cell line was chosen for its ability to form a brush border morphologically comparable to that of human colon, with heterogeneous microvillar presentation. Dendritic cells, derived by *in vitro* differentiation of primary human monocytes, were present on the basolateral face of the Transwell insert at approximately a 1:10 ratio to epithelial cells in the mature monolayer. Thus, each GIC unites a polarized human intestinal epithelial monolayer and dendritic cells, bringing together a mucus-bearing gut with essential elements of the innate immune system.

**Fig 2.**
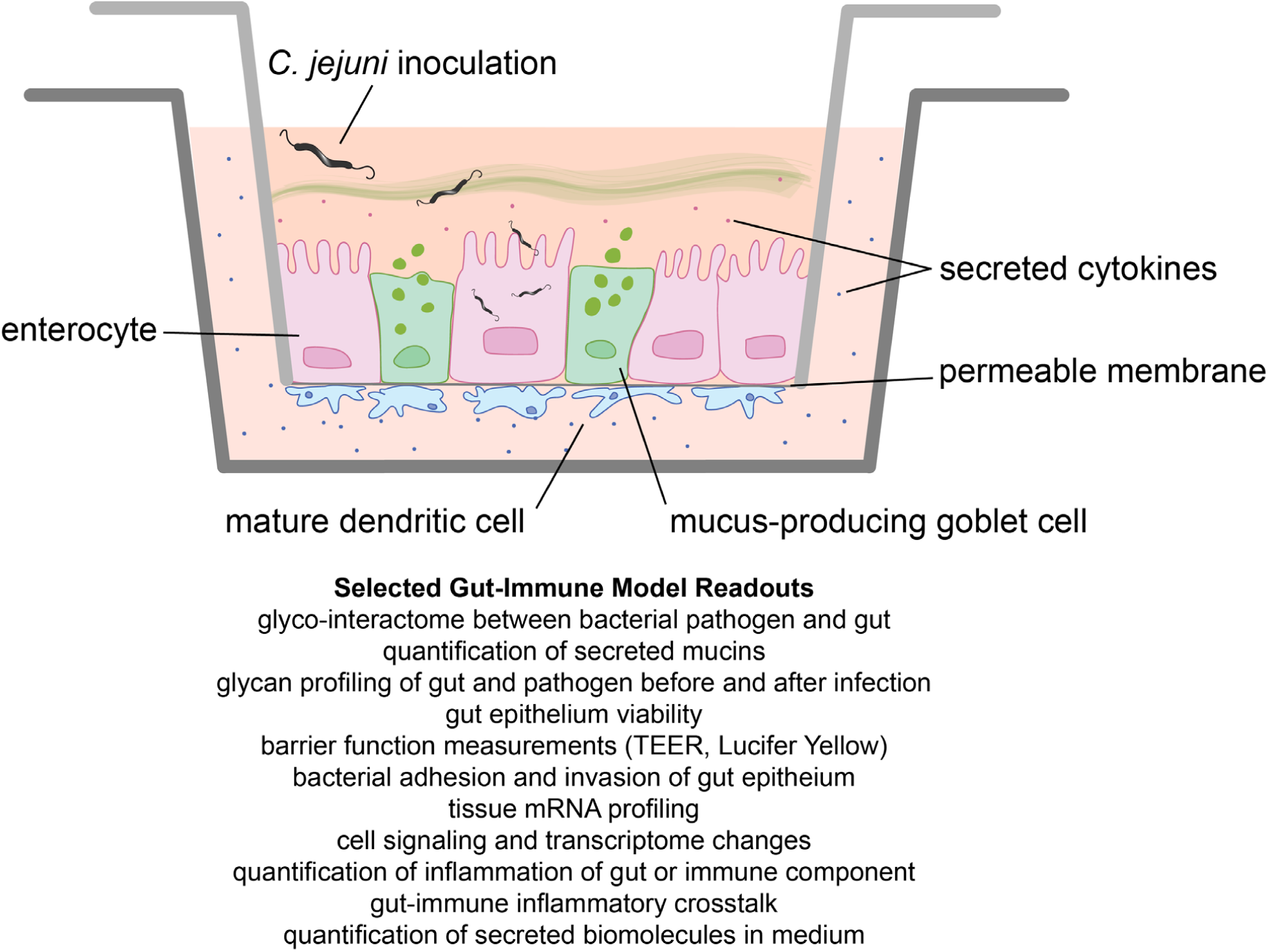
Schematic of the composition and readouts of gut-immune model within a Transwell insert.

This gut-immune co-culture model was used to quantify the impact of *C. jejuni* infection on different aspects of human intestinal function, such as barrier integrity, mucus production and innate immune system activation, and the importance of N-glycosylation in these host-pathogen interactions. We anticipate that the application of the gut-immune model to the characterization of this recalcitrant microaerophilic organism will advance studies of the pathogen in a relevant, quantitative multidimensional platform.

## Results

To demonstrate the utility of the gut-immune model for glycobiological studies of bacterial pathogenicity, the impact of *C. jejuni* N-glycosylation was characterized and quantified using several analytical readouts of the gut-immune model, listed in Figure 2. In these studies, the apical face of GICs was challenged with wildtype NCTC 11168 *C. jejuni* or a 11168Δ*pglE* strain (Linton et al. 2002), which exhibits near-complete abolition of N-linked protein glycosylation. (Fig S1).

It was initially unclear whether this fastidious microaerophilic organism, typically cultured to low optical densities in Mueller Hinton broth under 85% N_2_, 5% O_2_ and 10% CO_2_, would be viable or culturable during and after prolonged studies on GICs. Conversely, it was unknown if the nutrient-rich GIC growth media would cause bacterial overgrowth. However, culture conditions, multiplicities of infection and experimental time frames were identified that support bacterial and intestinal cell survival that would allow for measurements to be taken of both bacterial and mammalian processes.

Using these conditions, we characterized the impact of *C. jejuni* infection on aspects of GIC epithelial barrier integrity and probed the role of N-glycosylation. GICs were challenged with wildtype NCTC 11168 *C. jejuni* or 11168Δ*pglE* and intestinal secreted mucin content was examined. As shown in Figure 3A, the presence of both wildtype and Δ*pglE C. jejuni* resulted in a significant reduction of secreted mucins associated with the monolayer as compared with uninfected GICs. Next, changes in the barrier integrity in response to infection were quantified by Lucifer yellow permeability, measuring apical-to-basal transport of dye to indicate paracellular permeability in response to infection (Angelis and Turco 2011). In contrast to wildtype, which showed modest changes in barrier function, infection with Δ*pglE* resulted in significantly increased monolayer permeability (Fig 3B).

**Fig 3.**
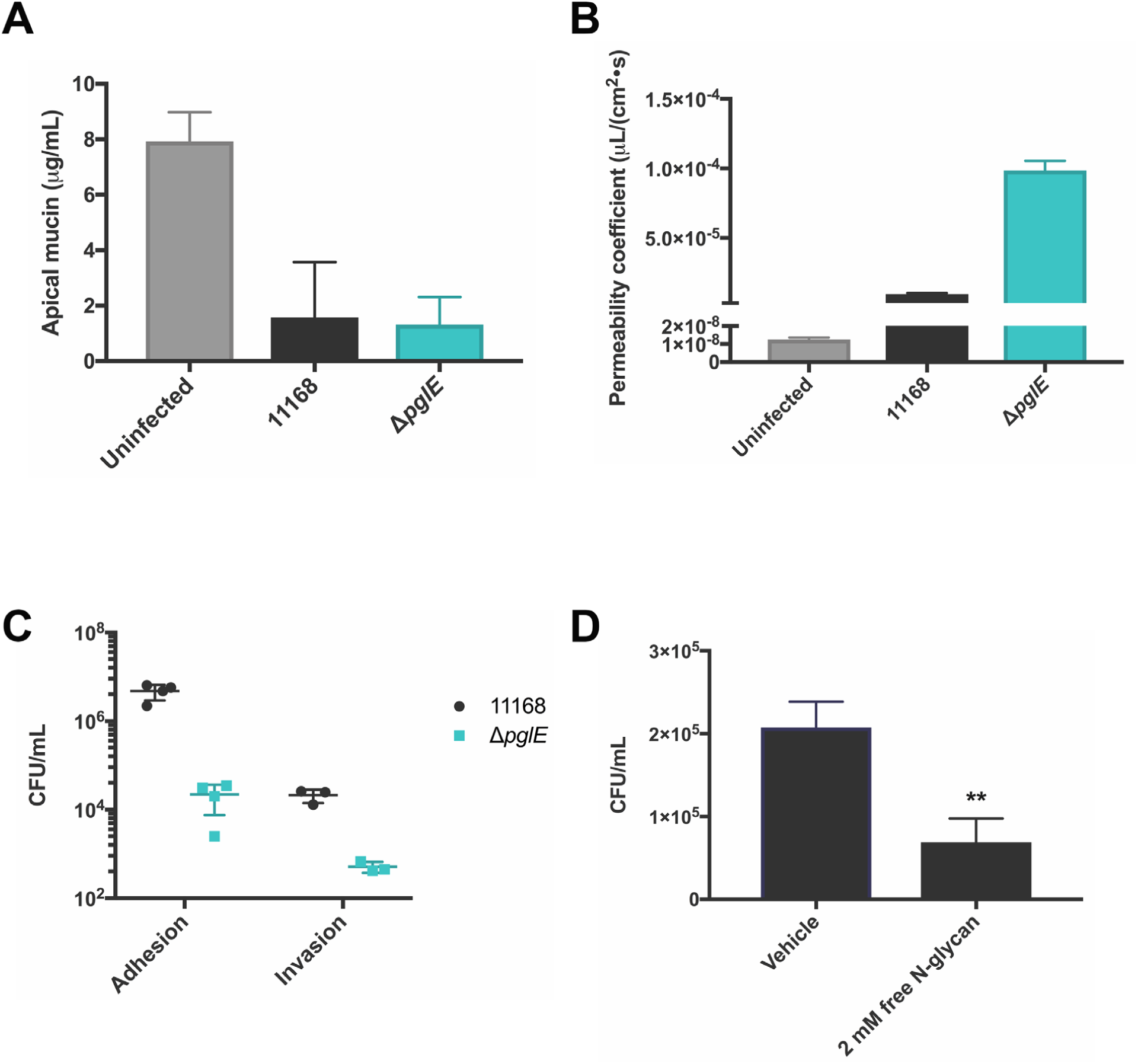
Effects of loss of N-glycosylation in 11168 *C. jejuni* on barrier functions and on epithelial adhesion and invasion. A) Infection of GICs with *C. jejuni* wildtype and Δ*pglE* results in significantly-reduced levels of soluble mucins in transwells. GICs were inoculated with 11168 or 11168Δ*pglE C. jejuni*, incubated for 24 h and soluble mucins quantified by Alcian blue colorimetric assay. Data shown are the average of three independent experiments, with error bars denoting one standard deviation. B) Glyco-deficient *C. jeuni* destabilize host tight junctions to a significantly higher degree than wildtype. GICs were inoculated with 11168 or 11168Δ*pglE C. jejuni* for 48 h and monolayer integrity characterized by Lucifer yellow permeability assay. Data shown are the average of 4 independent replicates, with error bars denoting one standard deviation. C) Loss of N-glycosylation in 11168 *C. jejuni* attenuates adhesion to and invasion of gut epithelial monolayer. GICs were inoculated with 11168 or 11168Δ*pglE C. jejuni* in DMEM growth media and incubated at 37 °C for 24 h. To measure adhesion, defined as bacteria adhered to or internalized into cells, monolayers were washed to remove non-adherent bacteria and lysed, followed by serial dilution and plating of lysates on *C. jejuni* selective media. To quantify internalized bacteria, monolayers were washed, subjected to gentamycin treatment and then lysed for plating on selective media. Data shown are independent biological replicates and representative of four independent data sets, with error bars denoting one standard deviation. Longer horizontal bar gives the mean of all measurements. D) Bacterial infectivity appears directly mediated by *C. jejuni* N-linked heptasaccharide. GICs were pre-incubated with fresh media (vehicle) or 2 mM free N-glycan, followed by 2 h incubation with 11168 *C. jejuni*. Data shown are the average of 3 independent replicates, in technical duplicate, with error bars denoting one standard deviation. ** indicates P<0.001 (Welch’s unpaired *t-*test)

Next, as pathogen/host association and invasion are primary hallmarks of *C. jejuni* infection *in vivo*, we quantified these behaviors in the GIC. In this analysis, a nearly 100-fold decrease in bacterial association with the epithelial monolayer between wildtype and Δ*pglE* was observed (Fig 3C). Monolayer infection was also quantified after gentamycin treatment of the apical compartment, killing all bacteria except those within cells, and a similar difference between wildtype and Δ*pglE* infectivity was observed. Additionally, a statistically-significant, but modest, 2.4-fold decrease in adhesion and invasion of wildtype *C. jejuni* was observed (Fig 3D) following treatment of monolayers with 2 mM free reducing heptasaccharide from *C. jejuni*, shown in Figure 1. Competition between soluble *C. jejuni* N-glycan, also known as free oligosaccharide or FOS (Dwivedi et al. 2013), and wildtype *C. jejuni* suggests a molecular role for the N-glycan in the adhesion and invasion of host intestinal cells.

A major advantage of the gut-immune model for studying *C. jejuni* pathogenicity is its utility in quantifying both intestinal epithelium inflammation and the resulting inflammatory response of the basal innate immune component. The extent of immunogenicity of *C. jejuni* infection was measured by apical infection of the GICs with wildtype or Δ*pglE C. jejuni* and sampling the basolateral media. Total cytokine and chemokine release into the basolateral compartment by both the intestinal epithelium and dendritic cells on the basal surface was measured. Several cytokines and chemokines were found to be increased in response to treatment with *C. jejuni*, with infection by Δ*pglE C. jejuni* resulting in slightly increased inflammation across many cytokines that were measured (Figs 4A, S2.) Together, these experiments show the utility of the GICs for studying the roles of protein glycosylation in the complex immune signaling resulting from the multiple cell types that compose human intestinal epithelia.

**Figure 4.**
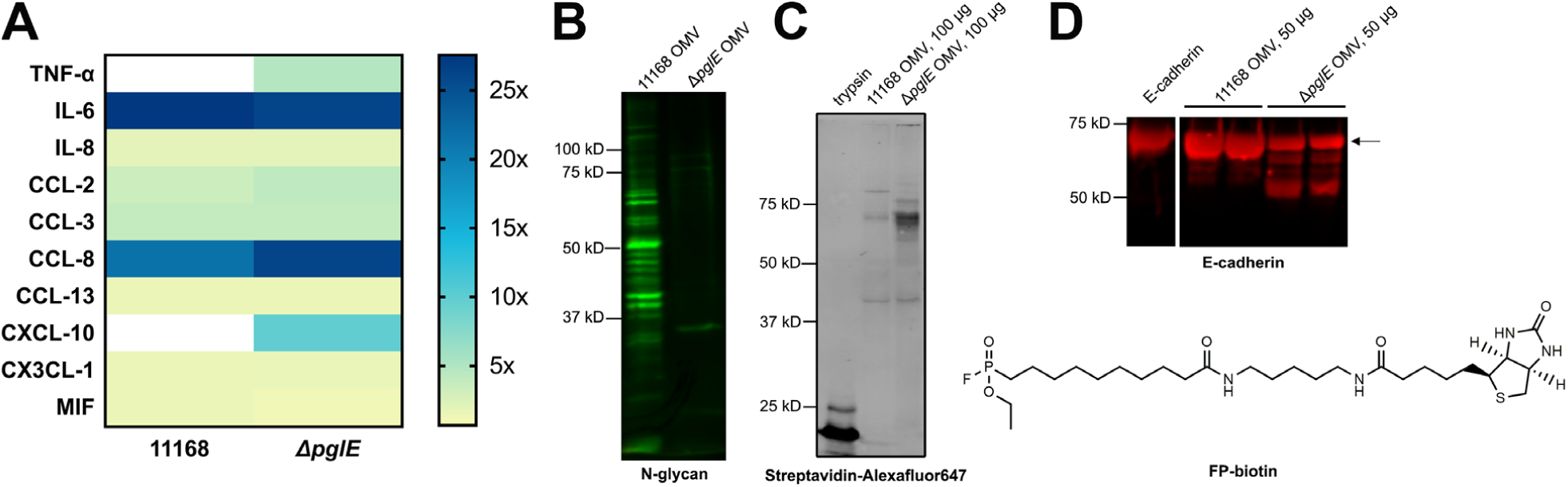
Loss of N-linked glycosylation in 11168 *C. jeuni* results in synergistic immune response on GICs and changes in virulence factor function. A) 11168 *C. jejuni* on apical face of gut-immune co-cultures elicit a synergistic immune response with dendritic cells present on the basolateral face. Growth media from GICs inoculated with 11168 *C. jejuni*, 11168Δ*pglE* or vehicle alone as a control were taken at 24 h post-inoculation from basolateral compartments. Inflammatory markers were quantified by 40-plex immunoassay. The average of four independent replicates, normalized to cytokine or chemokine concentrations in control GICs, are shown graphed as a heatmap showing fold-over vehicle alone. Cytokines and chemokine concentrations in all three data sets were compared to each other via 1-way ANOVA with no correction for multiple comparisons and those with an absolute P<0.05 shown. B) Loss of Pgl pathway function in 11168 *C. jejuni* results in the production of glycoprotein-deficient outer membrane vesicles. A 10 μg aliquot of OMV isolated from 11168 *C. jejuni* and 11168Δ*pglE* was taken up in loading dye, boiled, and visualized by immunoblotting with a rabbit anti-N-glycan antibody. C) Activity-Based Protein Profiling (ABPP) reveals OMV from 11168Δ*pglE C. jejuni* are enriched in serine proteases.A 100 μg aliquot of OMV isolated from 11168 *C. jejuni* and 11168Δ*pglE* was incubated for 30 min with FP-biotin reagent, followed by quenching of the labeling reaction with loading dye, followed by immunoblotting with Streptavidin-Alexafluor-647 and imaging. Trypsin was used as a positive control for ABPP. D) OMV from 11168Δ*pglE C. jejuni* cleave human tight junction proteins more readily compared to wildtype OMV. Recombinant His-tagged human E-cadherin was incubated for 16 h at RT with 50 μg OMV isolated from 11168 *C. jejuni* and 11168Δ*pglE*, followed by resolution by immunoblotting and probing with mouse anti-His antibody. Full length E-cadherin is indicated with an arrow.

Finally, to show the versatility of the GICs in glycobiological studies of secreted bacterial virulence factors, we characterized the effect of outer membrane vesicles (OMVs). OMVs are 25-250 nm diameter vesicles released by Gram-negative bacteria into the extracellular milieu. These bacterial “cargo drops” fuse with host cell membranes, delivering glycoproteins, effector proteins, DNA, endotoxins and proteases directly to the cytosol of host cells (Jan 2017, Schwechheimer and Kuehn 2015), initiating and facilitating pathogenic processes prior to direct cell-cell contact. OMVs from 11168 (Jang et al. 2014), hypermotile variant 11168H (Elmi et al. 2012) and 81-176 *C. jejuni* (Lindmark et al. 2009) have been shown to contain cytolethal distending toxin, and are hypothesized to be the primary mechanism of toxin delivery to the host. To study the role of the *pgl* locus on the behavior of this important *C. jejuni* virulence factor, OMVs were isolated from cultures of 11168 and 11168Δ*pglE C. jejuni* and analyzed for glycoprotein content by immunoblotting. A significant loss of N-linked glycoproteins was observed in OMVs isolated from Δ*pglE* when compared to wildtype (Fig 4B.) Analysis of wildtype and Δ*pglE* OMV content by activity-based serine protease profiling with the biotinylated fluorophosphonate FP-biotin (Fig 4C) (Kidd et al. 2001) detected an enrichment of proteases in Δ*pglE* OMV over those of wildtype 11168 *C. jejuni*. Proteases in OMVs from both strains were capable of cleaving human tight junction protein E-cadherin *in vitro*, with Δ*pglE* OMV resulting in more cleavage products (Fig. 4D).

A useful feature of GICs is the ability to vary composition of the apical epithelial monolayer according to investigator needs, while maintaining dendritic cells in the basolateral compartment. To study the functions of glycosylated and non-glycosylated *C. jejuni* OMV, barrier function and inflammation were measured on GICs lacking goblet cells and dendritic cells to ensure effective vesicle-epithelial cell contact. C2bbe1 monolayers were treated with 100 μg doses of OMVs from both strains and showed no change in barrier integrity by TEER (Fig. S3). However, IL-8, CXCL1, and several other immune markers were secreted by these GICs in response to OMV treatment (Fig S4). Importantly, these results illustrate the utility of GICs in quantitatively characterizing several aspects of virulence factor function such as those exhibited by *C.jejuni* OMV.

## Discussion and Conclusions

Use of the gut-immune model allows for convenient and simultaneous measurement of physiological and immunological responses to infection by *C. jejuni*, whose protein glycosylation pathway has been shown to be important in pathogenicity. We anticipate that this model system will be a valuable alternative for studying interactions of *C. jejuni* and other enteropathogens with host cells. These studies also show that GICs can be interfaced with microaerophilic pathogens such as *C. jejuni* for periods of time amenable to immunological study and under conventional culture conditions in normoxia and may be used to study other facets of this cell-cell interaction. Furthermore, the presented conditions should be readily adaptable to the study of other “difficult-to-culture” Gram-negative pathogens (e.g. *Helicobacter pylori*).

The analyses revealed some of the relationships between *C. jejuni* N-glycosylation and pathogenicity. Soluble mucin content on GICs (Fig 3A) was largely depleted during incubation with *C. jejuni* - a finding not previously quantified. The reduction of monolayer integrity (Fig 3B) was also directly correlated with loss of bacterial N-glycosylation and could be explained by a concomitant increase in proteases within Δ*pglE* OMVs capable of cleaving tight-junction proteins (Fig. 4D). Importantly, the degree of *C. jejuni* adhesion and invasion in the GICs directly correlated with global N-linked protein glycosylation and infectivity (Fig 3C).

Although direct release of cytokines by non-polarized and polarized epithelial monolayers and dendritic cells has been reported separately in response to wildtype *C. jejuni* exposure, this is the first characterization of the impact of the Pgl pathway on cytokine response in the intestinal epithelium in synergy with basolateral innate immune cells, with many cytokines and chemokines that were measured not previously reported in studies of *C. jejuni* infection. Here, it appears that basolateral inflammatory cytokine and chemokine secretion was modestly increased when the GICs were infected with the Δ*pglE* strain as compared to wildtype (Fig 4A). These cytokine profiles are likely dominated by factors produced by dendritic cells with some contribution from the epithelial cells; however, the exact contribution from each cell types is difficult to ascertain given the highly synergistic nature of the multicellular crosstalk. Finally, the glycosylation state of isolated wildtype and Δ*pglE* OMV did not change the inherent immunogenicity on C2bbe1-only GICs (Fig S4), and thus may not have a role in the loss of infectivity upon loss of glycosylation. However, it will be important to further characterize these virulence factors using other readouts provided by the gut-immune model.

In conclusion, the tripartite design of the gut-immune model, containing a recapitulated human intestinal epithelium and innate immune component, allowed for examination of the various roles played by the gut monolayer, the mucus, and dendritic cells and highlighted the importance of proper N*-*glycan presentation in various aspects of *C. jejuni* infectivity. This system also provided an opportunity to study the physiological and inflammatory interplay between all of the host components when exposed to this pathogen and its virulence factors – such insights would have otherwise been unobservable in other culture models. We anticipate this system will be advantageously applied to the study of other enteropathogens or aspects of pathogenic interactions.

## Acknowledgements

We thank Prof. Christine Szymanski for her valued advice and productive scientific discussions. The authors also thank the Szymanski laboratory at the Complex Carbohydrate Research Center (Athens, Georgia, USA) and Alberta Glycomics Centre (Edmonton, Alberta, Canada) for the generous gifts of the 11168Δ*pglE C. jejuni* strain, free oligosaccharide (FOS) and anti-N-glycan antibody. The authors also thank Prof. Ben Cravatt and his laboratory at Scripps Research Institute (La Jolla, CA, USA) for providing a sample of FP-biotin, and Dr. Enrique Valguarnera and Prof. Mario Feldman (Washington University School of Medicine, St. Louis, MO, USA) for technical advice on OMV preparation.

## Supplemental information

### Supplemental Figures

**Figure S1.**
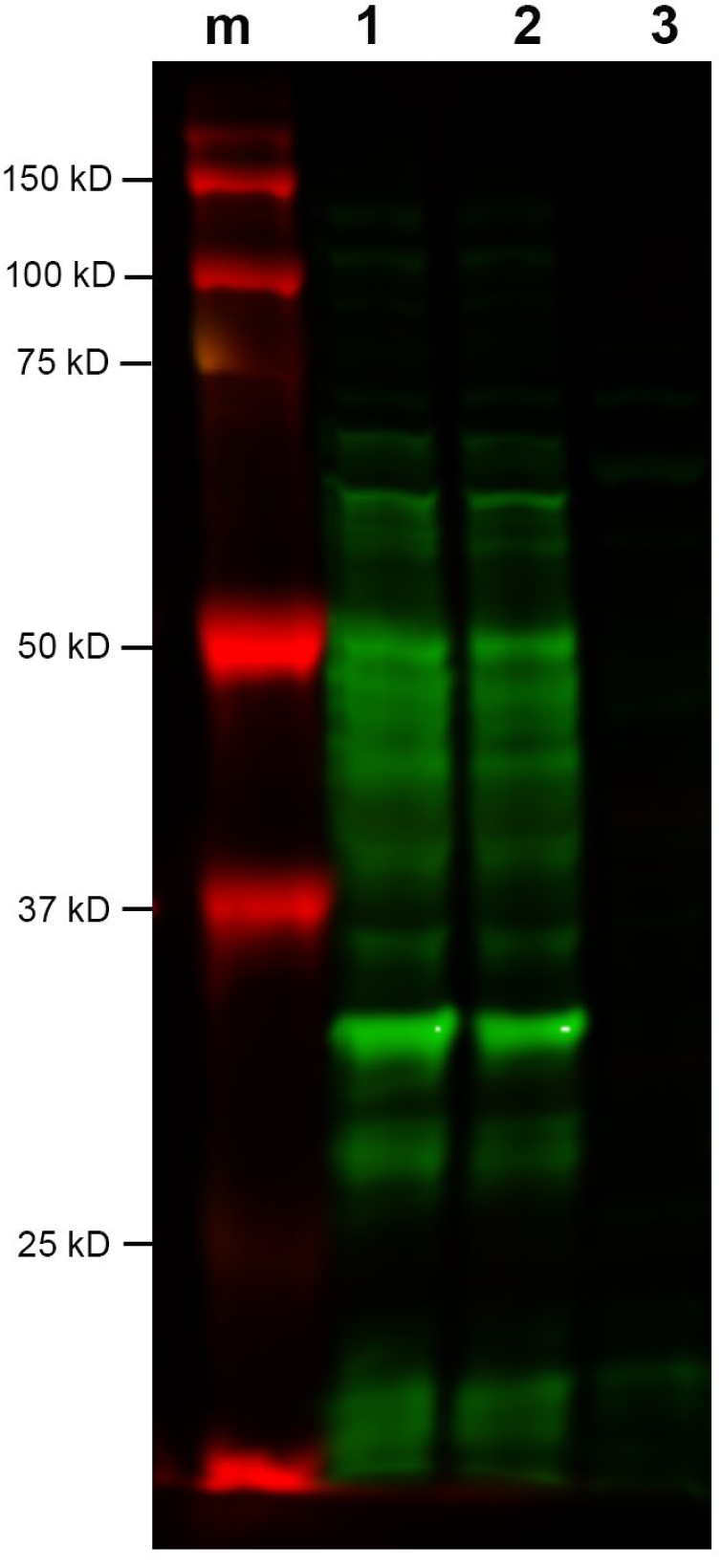
Knockout of *pglE* in 11168 *C. jejuni* results in significantly depleted *N*-glycosylated proteomes. Equal amounts of whole cell lysates of wildtype and Δ*pglE C. jejuni* were analyzed by immunoblotting, probing with rabbit anti-N-glycan antibodies. Secondary staining was performed with anti-rabbit Licor IrDye 800 antibodies and blots imaged using an Odyssey Infrared Imaging system. *Lanes*: m: molecular weight markers, 1: 11168 *C. jejuni*, 2: 11168, 3: 11168 Δ*pglE*. Lysate amounts were normalized by protein concentration, determined by Pierce BCA assay kit (Thermo Fisher Scientific, Waltham, MA.)

**Figure S2.**
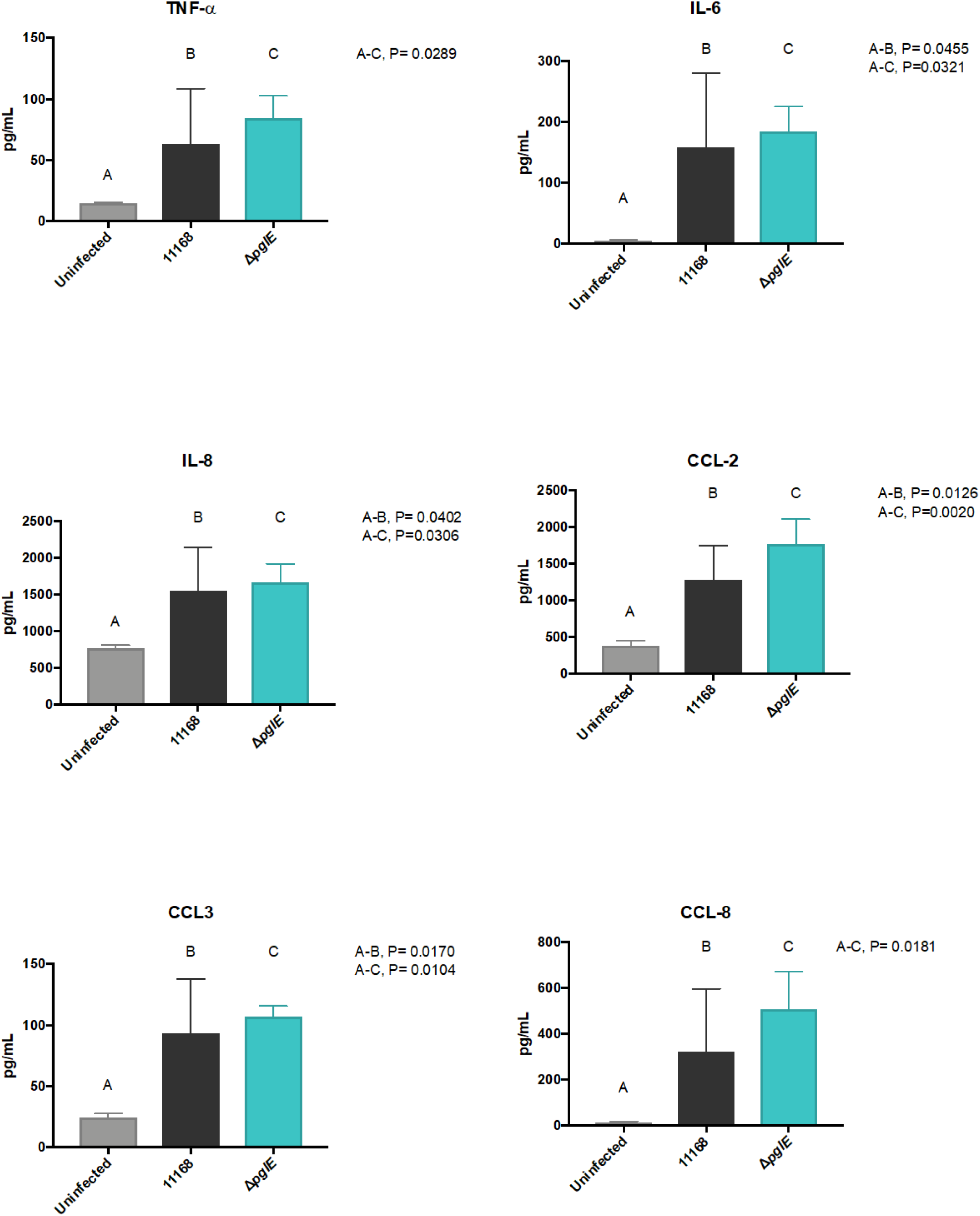
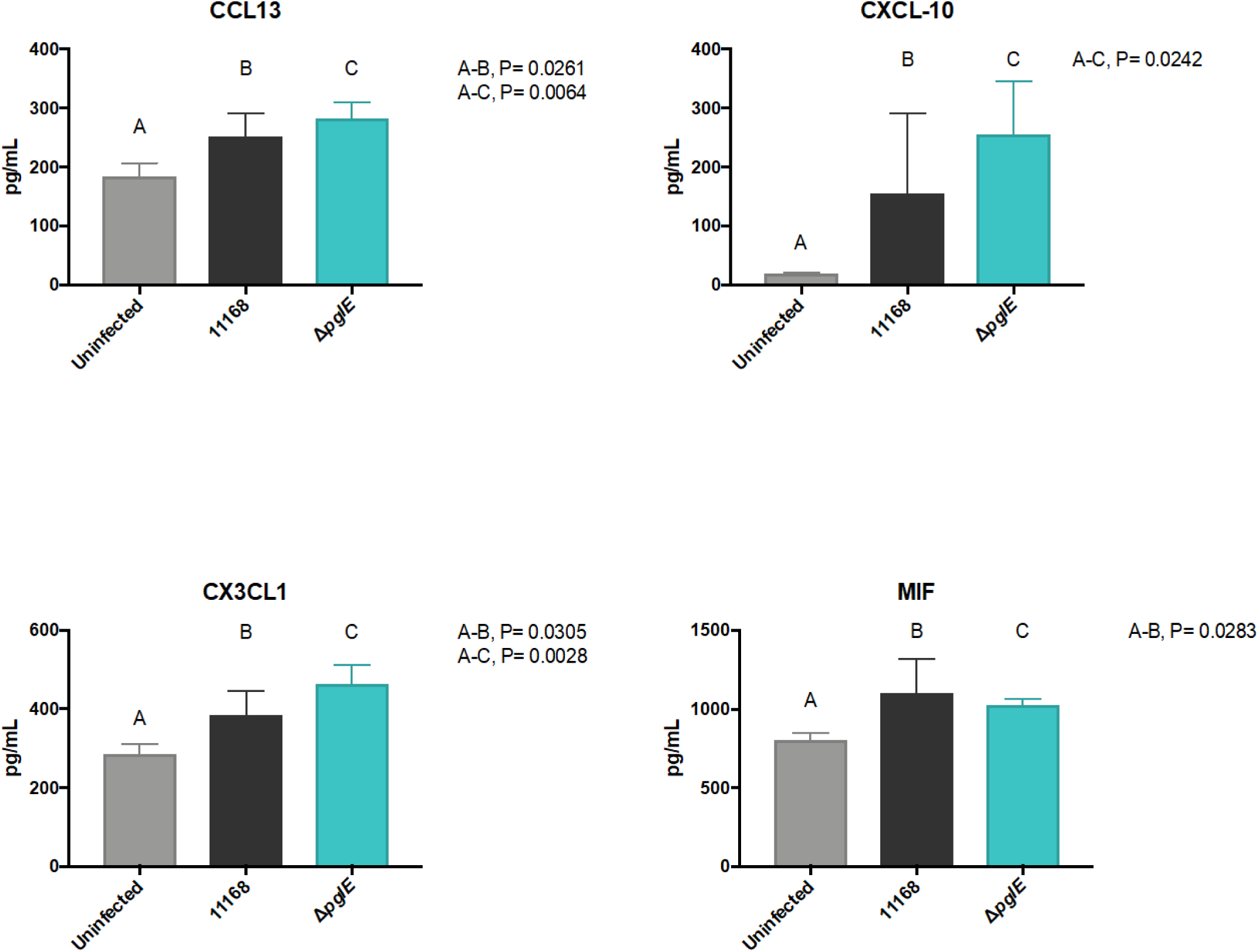
Treatment of GICs with *C. jejuni* results in synergistic release of several cytokines and chemokines into basal media by the epithelium and adjacent dendritic cells. GICs were inoculated with vehicle, 11168 *C. jejuni*, or 11168Δ*pglE* and incubated for 24 h. Basolateral media was sampled and inflammatory molecules quantified by 40-plex immunoassay. Bars show the mean of four independent replicates, with error bars showing one standard deviation. Mean marker concentrations in 11168 and 11168Δ*pglE* samples were compared to untreated gut MPSs and each other via 1-way ANOVA with no correction for multiple comparisons. P values for each statistically significant comparison are shown in each legend.

**Figure S3.**
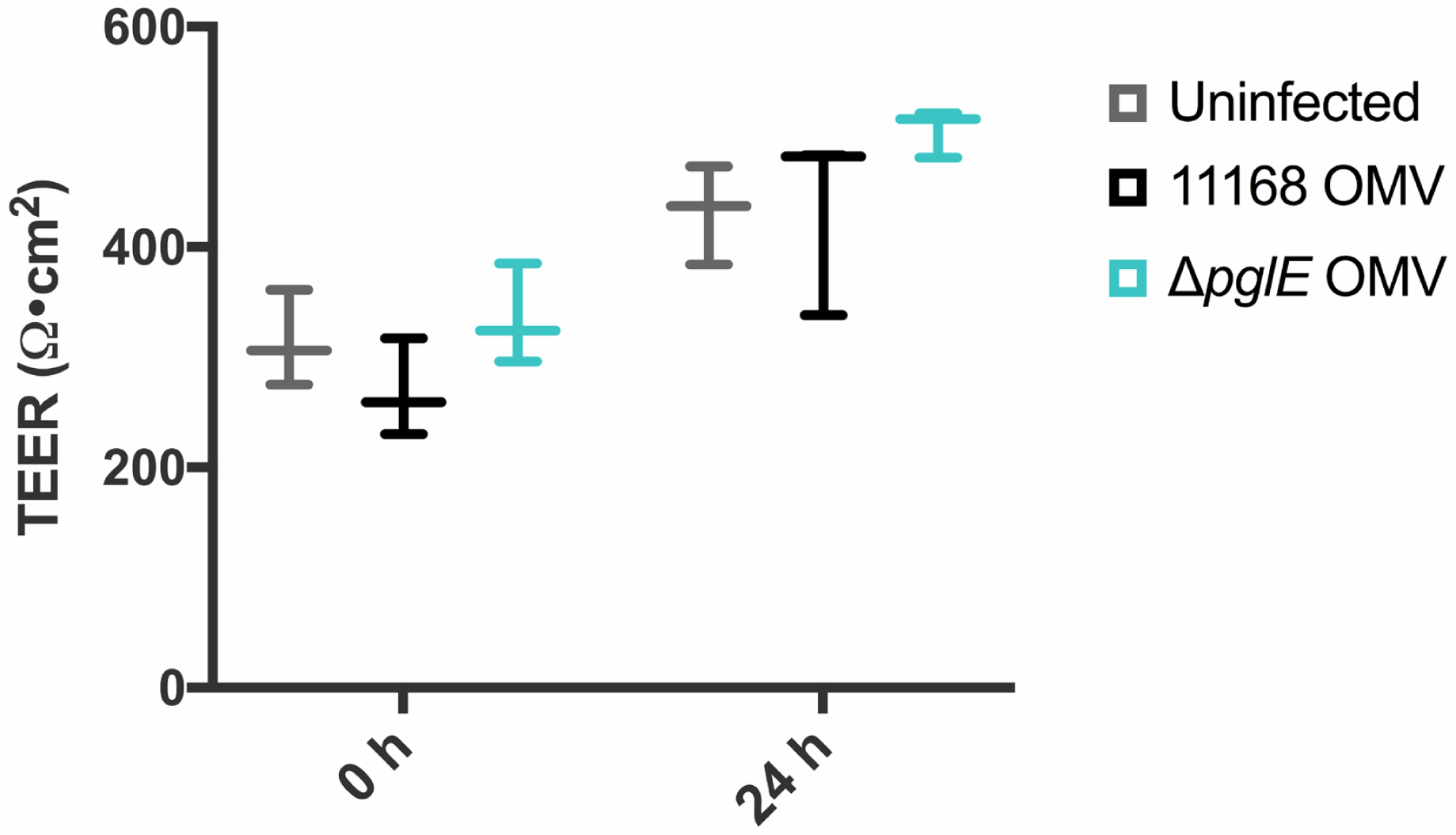
OMV from 11168 *C. jejuni* and variant do not provoke measurable changes in barrier integrity by TEER. Barrier integrity of each C2BBe1-only GIC was quantified by transepithelial electrical resistance (TEER) measurements at the start of the experiment. GICs were treated with vehicle, 100 μg OMV isolated from 11168 *C. jejuni*, or 100 μg 11168Δ*pglE* OMV and incubated at 37 °C for 24 h. After incubation, TEER values were again measured for all monolayers and the apical media was sampled for immunoassay analysis. Data shown are the average of three independent replicates, with error bars denoting one standard deviation. The mean is shown by a longer horizontal bar.

**Fig S4.**
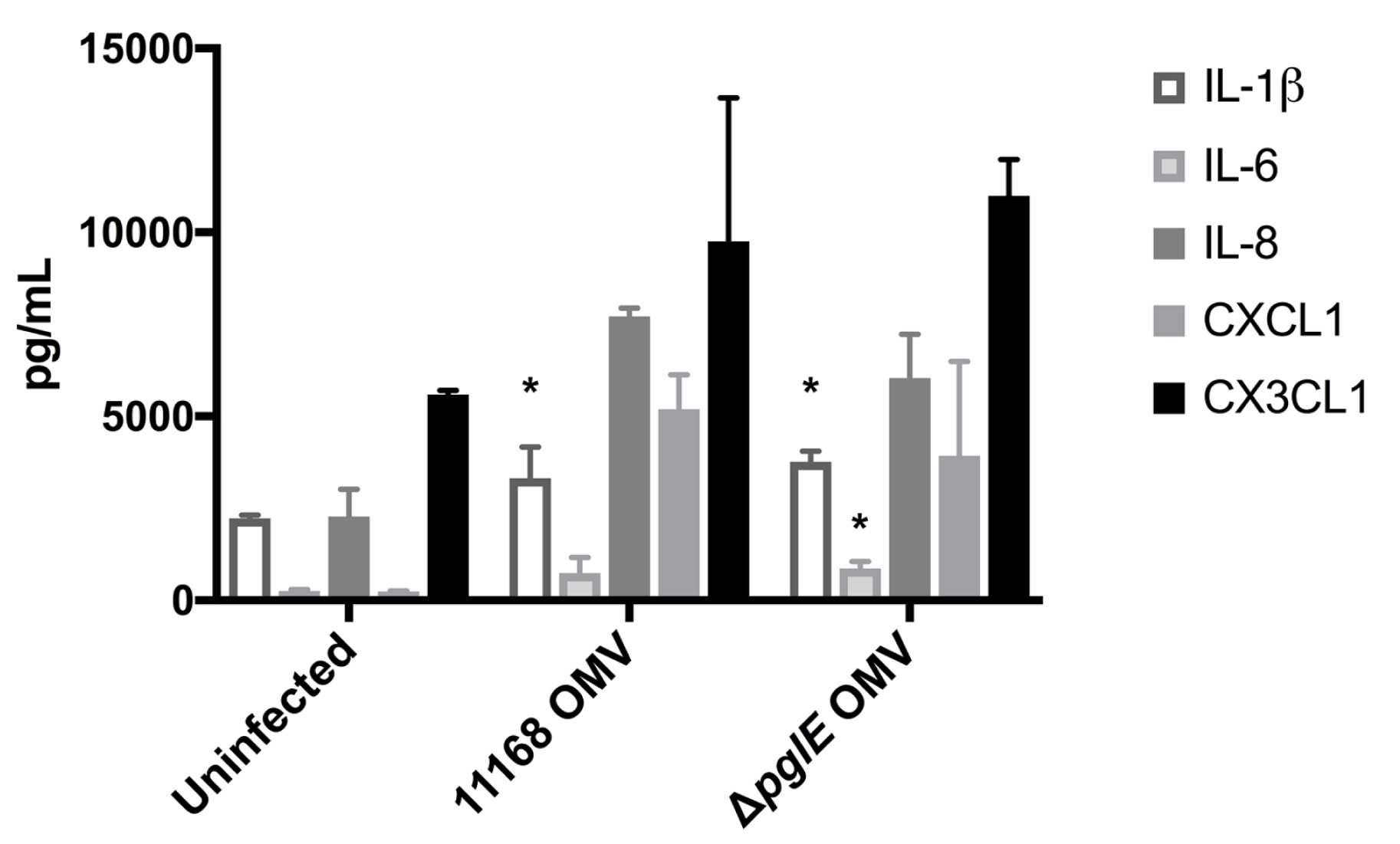
OMV from 11168 *C. jejuni* provoke an immune response from gut epithelia. GICs were treated with vehicle alone, 100 μg OMV isolated from 11168 *C. jejuni*, or 100 μg 11168Δ*pglE* OMV, followed by incubation for 24 h. Apical media from all GICs was sampled and cytokines and chemokines present quantified by multiplexed immunoassay. Data shown are the average of three independent replicates, in technical duplicate, with error bars denoting one standard deviation. Mean marker concentrations in 11168 OMV and 11168Δ*pglE* OMV samples were compared to uninfected GICs and each other via 1-way ANOVA with no correction for multiple comparisons. * indicates statistically significant differences with absolute P-values ≤0.05, ** indicates P≤0.01, *** indicates P≤0.001.

## Materials and Methods

### General information

All common chemicals, gases and reagents were purchased from Sigma-Aldrich, AirGas, Fisher Scientific and VWR unless otherwise noted. Following is a list of the sources of other key reagents and expendable materials used in these studies: BBL Mueller-Hinton broth (BD and Co., Cat. No. 211443); Trimethoprim (Chem Impex, Cat. No. 01634); kanamycin sulfate (Teknova, Cat. No. K2151); Oxoid AGS plastic pouches (Invitro Diagnostics, Cat. No., AG0020C); sealing clips for plastic pouches (Invitro Diagnostics, Cat. No. AN0005C); culture tubes for bacterial cultures (17 × 100 mm) (VWR, Cat. No. 60818-689); Pierce BCA assay kit (Thermo Fisher Scientific, Cat. No. 23227); mouse anti-6x-His epitope tag monoclonal antibody (Thermo Fisher Scientific, Cat. No. MA1-21315); LI-COR IRDye 800CW goat anti-rabbit IgG (LI-COR, Cat. No. 926-32211); Licor IRDye 680LT goat anti-mouse IgG (LI-COR, Cat. No. 680 926-68020); LI-COR IRDye 800CW goat anti-mouse IgG (LI-COR, Cat. No. 926-32210); streptavidin-AlexaFluor 647 (Thermo Fisher Scientific, Cat. No. S32357); 75 cm^2^ tissue culture flasks (Corning, Cat. No. 353136); trypsin-EDTA (Gibco, Cat. No. 252000-56); Transwell membrane inserts (Corning, Cat. No. 3460); trypsin (Corning, Cat. No. 25-051-CI); recombinant E-cadherin (R & D Systems, Cat. No. 8505-EC-050).

### Tissue culture media compositions

Gut Cell Line Media: DMEM (Gibco, Cat. No. 11965), 10% fetal bovine serum (FBS, Atlanta Biologicals, Cat. No. S11150), 1x non-essential amino acids (NEAA, Gibco, Cat. No. 11140-148), 1x GlutaMAX (Gibco, Cat. No. 35050-061), 1% penicillin-streptomycin (P/S, Gibco, Cat. No. 15140-148.)

Gut Insert Seeding Media: Advanced DMEM (Gibco, Cat. No. 12491), 10% FBS, 1x GlutaMAX, 1% P/S.

Serum-Free Insert Medium: Advanced DMEM, 1x insulin-transferrin-selenium supplement (ITS, Roche, Cat. No. 11074547001), 1x GlutaMAX, 1% P/S.

Apical Media: DMEM, ITS, 1x GlutaMAX, 1x NEAA and 1% P/S.

Basal Media: Advanced DMEM, 1x ITS, 1x GlutaMAX, 0.85-1.25 mg/mL bovine serum albumin (BSA, Sigma-Aldrich, Cat. No. A9576) and 1% P/S.

### Generalized Procedure A: Culture of *C. jejuni* strains

Samples from glycerol stocks of *C. jejuni* (NCTC 11168 or 11168Δ*pglE*) were streaked onto MH agar plates containing selection antibiotics (10 μg/mL trimethoprim for 11168 wildtype, 10 μg/mL trimethoprim and 20 μg/mL kanamycin sulfate for 11168Δ*pglE*). Plates were placed inside a plastic pouch (along with an atmospheric CO_2_ indicator) and purged with a microaerophilic gas mixture (85% N_2_, 10% CO_2_, 5% O_2_) multiple times until the correct concentration of CO_2_ was reached. Sealed pouches were then incubated at 37 °C for 24 h, after which colonies were taken up in 3 mL of PBS broth. Cells taken from these master stocks of *C. jejuni* were utilized in the subsequent experiments.

### Generalized Procedure B: Maintenance and passaging of intestinal epithelial cell lines C2BBe1 and HT29-MTX

C2BBe1 (ATCC, passage 48-58) and HT29-MTX (Sigma, passage 20-30) were maintained with Gut Cell Line Media in separate 75 cm^2^ flasks. Cells at 80-90% confluence were passaged by washing with PBS without calcium and magnesium (PBS-/-) and lifting with 0.25% trypsin-EDTA at 37°C. After 10 minutes, cells were manually dissociated, quenched with media, and centrifuged at 300 *x g* for 5 min in 15 mL conical tubes. After, cells were re-suspended and quantified by Trypan Blue exclusion for continued passaging or seeding into Transwell membrane inserts according to *Generalized Procedures C* and *D*. Passages of C2BBe1 and HT29-MTX were seeded with 6 ×10^5^ and 2 ×10^6^ cells, respectively, every 6-7 days and were fed with Gut Cell Line Medium every 2-3 days.

### Generalized Procedure C: Preparation and maintenance of polarized epithelial monolayers

Polarized epithelial monolayers were prepared as previously described (Chen et al. 2017). Prior to seeding, Transwell membrane inserts were coated with 50 mg/mL rat tail collagen I (Corning, Cat No. 354236) in PBS-/- for 2 h at rt. Intestinal epithelial cells were collected according *Generalized Procedure B*, ensuring cells had been passaged twice post-thawing prior to use. For C2BBe1-only monolayers, after rinsing away collagen coating solution with Advanced DMEM, 1.12 × 10^5^ C2BBe1 cells (10^5^ cells/cm^2^) were seeded in each insert in 500 μL of Gut Seeding Media to form C2BBe1 monolayers. For C2BBe1/HT29-MTX monolayers, cells seeded on inserts were a 9:1 mixture of the C2BBe1 and HT29-MTX cell lines. All inserts were fed every 2-3 days with 500 μL and 1.5 mL Gut Insert Seeding media in apical and basolateral chambers, respectively. From Day 7 on, inserts were fed every 2-3 days with Serum-Free Insert Medium. Inserts were cultured in this manner for 21-27 days before use in any experiments.

### Generalized Procedure D: Immune cell isolation, differentiation and seeding

Peripheral blood mononuclear cells (PBMCs) were processed from Leukopak (STEMCELL Technologies, Cat. No. 70500) using the recommended protocol for cell isolation without density centrifugation. Isolated cells were aliquoted and stored in liquid nitrogen.

To derive dendritic cells, monocytes were isolated from thawed PBMCs using the EasySep Human Monocyte Enrichment Kit without CD16 Depletion (STEMCELL Technologies, Cat. No. 19058). The purified monocytes were seeded into a 24-well tissue culture treated plate at ~1.2 million cells/well in Advanced RPMI medium (Gibco, Cat. No. 12633-012) supplemented with 1X GlutaMax, 1% P/S, 50 ng/mL GM-CSF (Biolegend, Cat. No. 572903), 35 ng/mL IL-4 (Biolegend Cat. No. 574004) and 10 nM retinoic acid. After 7 days of differentiation (at day 19-20 of the epithelial culture), mature dendritic cells were lifted using PBS-/- and Accutase (Gibco, Cat. No. A11105-01) and seeded onto the basolateral side of inverted gut Transwell inserts at 10^5^/insert, allowing 2 h for attachment.

### Preparation of retinoic acid stock solution

Retinoic acid was dissolved in 200 proof EtOH and concentration was determined by measuring absorbance at 350 nm (ε_max_ (EtOH) = 44,300 M^−1^cm^−1^). The solution was diluted to 50μM in PBS-/- with 1% BSA. Frozen stocks were kept at −20°C for up to 3 months.

### Generalized Procedure E: Preparation of gut-immune co-cultures (GICs)

To begin, intestinal epithelia were prepared as described in *Generalized Procedure C*. At Day 13, monocytes were isolated and differentiated as in *Generalized Procedure D*. On Day 20 post-monolayer seeding (7 days post-monocyte isolation), dendritic cells were recovered and seeded onto the basal membrane of the gut cultures. From this point, gut-immune co-cultures were maintained with 500μL Apical Media and 1.5 mL Basal Media in their respective chambers. GICs were utilized in experiments 21 days post-monolayer seeding. GICs were rinsed with Serum-Free Maintenance Media without antibiotics prior to inoculation with *C. jejuni*.

### Generalized Procedure F: Treatment of GICs with strains and mutants of *C. jejuni*

The apical compartment of GICs containing gut epithelial monolayers or co-cultures were inoculated with *C. jejuni* strains and mutants ahead of experiments characterizing bacterial adhesion and internalization under various conditions and ahead of immunoassays and other measures of epithelial health. To begin, *C. jejuni* stocks were prepared according to General Procedure A and aliquots sufficient to give an OD = 1.0 in 1 mL were taken into microcentrifuge tubes. Cells were centrifuged at 16,000 *x g* for two minutes to give a pellet (pink in color,) the supernatant removed and the pellet resuspended in 1 mL DPBS. Apical growth media without P/S was warmed to 37 °C and inoculated with washed *C. jejuni* to give a solution with initial OD of 0.03 (multiplicity of infection: 10) or 0.15 (multiplicity of infection: 50). GICs to be treated were provided fresh basal growth media right before innoculation. Growth media was aspirated from the apical face of each GIC to be treated and replaced with 500 μL of media containing *C. jejuni* strains.

### Measurement of bacterial adhesion adhesion to GIC epithelia

GICs inoculated with wildtype *C. jejuni* NCTC 11168 or 11168Δ*pglE* according to General Procedure F were incubated at 37 °C for 24 h, after which inserts were visually inspected and apical compartments washed four times with 500 μL DPBS to remove non-adherent bacteria. GICs were given 500 μL DPBS with 0.1% Triton and placed on an orbital shaker shaking at 300 rpm for 20 min to lyse epithelial monolayers. Lysates containing bacteria from monolayer surfaces and cell interiors were serially diluted in MH broth and 50 μL plated on MH agar containing appropriate selection antibiotics for colony counting.

### Measurement of invasion by C. jejuni into epithelial monolayers by gentamycin treatment

GICs inoculated with wildtype *C. jejuni* NCTC 11168, or 11168Δ*pglE* according to General Procedure C were incubated at 37 °C for 24 h, after which inserts were visually inspected and monolayer washed 3 times with 500 μL DPBS to remove non-adherent bacteria. GICs were then treated with 500 μL apical media containing 210 μM gentamycin sulfate and incubated at 37 °C for 45 min. GICs were washed four times with 500 μL DPBS to remove bacteria and residual antibiotic, followed by treatment with 500 μL DPBS with 0.1% Triton. GICs were placed on an orbital shaker shaking at 300 rpm for 20 min to lyse and lysates were serially diluted in MH broth and plated on MH agar containing appropriate selection antibiotics for colony counting.

### Secreted mucin quantification by Alcian blue colorimetric assay

Secreted mucins in apical media collected from experiments were quantified *via* a colorimetric assay adapted from Hall et al (Hall Rl Fau - Miller et al.). Briefly, collected samples stored in low-binding microcentrifuge tubes were spun at 6,000 *x g* for 5 min to pellet bacteria and/or cell debris prior to analysis. Supernatants were analyzed immediately or flash frozen with liquid nitrogen for storage at −80 °C. Samples were mixed with Alcian Blue solution (Richard Allan Scientific Co., San Diego, CA) in a ratio of 3:1 and incubated for 2 h. After, samples in 96-well plates were centrifuged at 1,640 *x g* for 30 min at rt. Supernatants were removed by inversion and the resulting pellet resuspended twice with a 40% ethanol/60% 0.1M sodium acetate buffer with 25 mM MgCl_2_, pH 5.8, centrifuging 10 min as above after each resuspension. Washed pellets were fully dissolved in 10% SDS solution and absorption was measured at 620 nm on a plate reader (Spectramax m3/m2e). Mucin from bovine submaxillary glands (Sigma, Cat. No. M3895) served as a standard and was prepared and analyzed identically in parallel with experimental samples.

### Fluorometric quantification of paracellular permeability in epithelial monolayers

Paracellular permeability of epithelial monolayers was quantified utilizing Lucifer Yellow reagent according to manufacturer’s instructions. The epithelial monolayers of GICs were washed with transport buffer (HBSS with CaCl_2_, MgCl_2_ supplemented with 10 mM HEPES, pH 7.4) and 500 μl of 100 mM Lucifer Yellow in transport buffer was added to the apical compartment of each insert. Transport buffer (1.5 mL) was added to the basal compartment and inserts were incubated at 37 °C in 5% CO_2_ for 1-2 h with shaking. Following incubation, inserts were removed and 150 μl of each sample transferred to a 96-well plate and fluorescence measured (λ_ex_= 485 nm, λ_em_= 530 nm.) Standard curves of Lucifer Yellow alone in transport buffer were generated to determine fluorophore concentrations in experimental samples. Apparent permeability coefficients were calculated according to manufacturer’s instructions.

### Quantification of cytokine/chemokine release by multiplexed immunoassay

Cytokine/chemokine amounts in apical or basal media samples collected from experiments were measured using Bio-Plex Pro Human Chemokine Panel, 40-plex (Bio-Rad Laboratories, Inc., Cat. No. 171AK99MR2) according to manufacturer’s instructions. Collected samples stored in low-binding microcentrifuge tubes were spun at 6,000 *x g* for 5 min to pellet bacteria and/or cell debris prior to analysis. Supernatants were analyzed immediately or flash frozen with liquid nitrogen for storage at −80 °C. BSA was added to all samples (final concentration 5 mg/mL) in order to minimize non-specific binding to beads. Multiple dilutions of each sample were analyzed to ensure to ensure measurements were within the linear dynamic range of the assay. Assays were performed on a Bio-Plex 3D Suspension Array System and data collected using xPONENT for FLEXMAP 3D software, version 4.2 (Luminex Corporation, Austin, TX). Concentrations of analytes were determined utilizing a standard curve generated by fitting a 5-parameter logistic regression of mean fluorescence on known concentrations of each analyte with Bio-Plex Manager software.

### Isolation of outer membrane vesicles from NCTC 11168 and 11168 Δ*pglE*

NCTC 11168 and NCTC 11168Δ*pglE* from 24 h plates were used to inoculate 1 L of Mueller Hinton media. Cultures were grown overnight under microaerophilic conditions at 37 °C with shaking to mid log phase. OMV isolation was adapted from Valguarnera et al (Valguarnera and Feldman 2017). Briefly, bacteria were pelleted for 30 min at 3,600 *x g* and the supernatant filtered through 0.22 μm membrane. The filtrate was ultracentrifuged for 2 h at 100,000 *<u>x g</u>*. The vesicles were resuspended in PBS and ultracentrifuged an additional 2 h at 100,000 *x g*. Fresh PBS was added and the vesicles were ultracentrifuged overnight at 200,000 *x g*. The supernatant was removed and the vesicles were brought up in 2 ml in PBS and the protein content was assessed by BCA assay.

### Characterization of *C. jejuni* OMV by SDS-PAGE and immunoblotting

For analysis of total protein content, 100 μg of purified OMV was mixed with 20 μl 2x Laemmli buffer and boiled for 10 min. The samples were separated by 10% SDS-PAGE and stained with Instant Blue overnight. For N-glycan analysis, 20 μg of the OMV was separated by 10% SDS PAGE, transferred to nitrocellulose membrane by wet transfer, and blocked with 5% BSA in TBS for 1 h. The membrane was blotted with rabbit anti-N-glycan antibody (1:10,000 dilution) overnight at 4 °C, followed by secondary anti-rabbit antibody incubation 1:10,000 dilution for 2 h. Blots were visualized by LI-COR Odyssey scanner.

### Serine protease profiling of *C. jejuni* OMV with FP-biotin

Active serine protease profiling was performed using 100 μg purified OMV as previously described (Liu et al. 1999). Samples were incubated with 2 μM **FP-biotin** (10-(fluoroethoxyphosphinyl)-*N*-(biotinamidopentyl)decanamide) in a final volume of 100 μl in PBS pH 7.3 for 30 minutes at room temperature. Four μg of trypsin was included as a positive control for serine protease activity. The samples were evaporated to dryness and reconstituted in 20 μl 1x Laemmli buffer and boiled for 10 min. The samples were separated by SDS PAGE (10% polyacrylamide gels) and transferred to nitrocellulose membrane by wet transfer method. The blot was blocked overnight in 10% wt/vol dry milk in TBS then probed with streptavidin-Alexafluor 647 at a 1:1,000 dilution for one h, and visualized with LI-COR Odyssey scanner.

### E-cadherin cleavage assay with *C. jejuni* OMV

The cleavage of recombinant E-cadherin was determined as described previously(Elmi et al. 2016). Briefly, 10 μg of purified OMV was incubated with 1 μg of recombinant human E-cadherin containing a C-terminal 6xHis tag in PBS at 37 °C for 16 h. The reactions were mixed 1:1 with 2 x Laemmli buffer and boiled for 10 minutes. The samples were separated by SDS PAGE and transferred to nitrocellulose membrane by wet transfer. The membrane was blocked with 5% BSA in TBS for one h then incubated with mouse anti-His antibody overnight at 4 °C. The blot was incubated with anti-mouse secondary antibody at a 1:10,000 dilution for one h then imaged with LI-COR Odyssey scanner.

### Barrier function characterization via transepithelial electrical resistance (TEER)

TEER measurement across epithelial monolayers in GIC transwells was carried out using EndOhm-12 chambers and an EVOM2 meter (World Precision Instruments, Sarasota, FL). Transwells and EndOhm chamber were maintained at 37 °C during all measurements to minimize variability. TEER of C2BBe1-only GICs in transwells (Day 21) was measured at the start of the experiment. Next, the apical media of the transwells was aspirated and replaced with 500 μL apical media alone or 500 μL containing 10 or 100 μg of wildtype or Δ*pglE* OMV. Transwells were incubated at 37 °C for 24 h, after which apical media was sampled and TEER measured a second time. Sampled media was flash frozen in liquid nitrogen for immunoassay analysis.

### Statistical Analysis

All experiments were performed at least in triplicate, with additional replicates, technical replicates and independent biological replicates noted in each figure caption. Statistical analyses of data sets were performed using GraphPad Prism software. Statistical significance (P < 0.05) and P values were calculated utilizing Student’s t-tests or 1-way ANOVA as appropriate.

## References

Global priority list of antibiotic-resistent bacteria to guide research, discovery, and development of new antibiotics; world health organization (who): Geneva, 2017.

Alemka A, Clyne M, Shanahan F, Tompkins T, Corcionivoschi N, Bourke B. 2010. Probiotic colonization of the adherent mucus layer of ht29mtxe12 cells attenuates <em>campylobacter jejuni</em> virulence properties. Infection and Immunity, 78:2812–2822.

Alemka A, Corcionivoschi N, Bourke B. 2012. Defense and adaptation: The complex inter-relationship between campylobacter jejuni and mucus. Front Cell Infect Microbiol, 2:15.

Angelis ID, Turco L. 2011. Caco-2 cells as a model for intestinal absorption. Current Protocols in Toxicology, 47:20.26.21–20.26.15.

Backert S, Boehm M, Wessler S, Tegtmeyer N. 2013. Transmigration route of campylobacter jejuni across polarized intestinal epithelial cells: Paracellular, transcellular or both? Cell Commun Signal, 11:72.

Bahrami B, Macfarlane S, Macfarlane GT. 2011. Induction of cytokine formation by human intestinal bacteria in gut epithelial cell lines. J Appl Microbiol, 110:353–363.

Chen WLK, Edington C, Suter E, Yu J, Velazquez JJ, Velazquez JG, Shockley M, Large EM, Venkataramanan R, Hughes DJ, Stokes CL, Trumper DL, Carrier RL, Cirit M, Griffith LG, Lauffenburger DA. 2017. Integrated gut/liver microphysiological systems elucidates inflammatory inter-tissue crosstalk. Biotechnol Bioeng, 114:2648–2659.

Dorrell N, Mangan JA, Laing KG, Hinds J, Linton D, Al-Ghusein H, Barrell BG, Parkhill J, Stoker NG, Karlyshev AV, Butcher PD, Wren BW. 2001. Whole genome comparison of campylobacter jejuni human isolates using a low-cost microarray reveals extensive genetic diversity. Genome Res, 11:1706–1715.

Dwivedi R, Nothaft H, Reiz B, Whittal RM, Szymanski CM. 2013. Generation of free oligosaccharides from bacterial protein n-linked glycosylation systems. Biopolymers, 99:772–783.

Edington CD, Chen WLK, Geishecker E, Kassis T, Soenksen LR, Bhushan BM, Freake D, Kirschner J, Maass C, Tsamandouras N, Valdez J, Cook CD, Parent T, Snyder S, Yu J, Suter E, Shockley M, Velazquez J, Velazquez JJ, Stockdale L, Papps JP, Lee I, Vann N, Gamboa M, LaBarge ME, Zhong Z, Wang X, Boyer LA, Lauffenburger DA, Carrier RL, Communal C, Tannenbaum SR, Stokes CL, Hughes DJ, Rohatgi G, Trumper DL, Cirit M, Griffith LG. 2018. Interconnected microphysiological systems for quantitative biology and pharmacology studies. Sci Rep, 8:4530.

Elmi A, Watson E, Sandu P, Gundogdu O, Mills DC, Inglis NF, Manson E, Imrie L, Bajaj-Elliott M, Wren BW, Smith DGE, Dorrell N. 2012. Campylobacter jejuni outer membrane vesicles play an important role in bacterial interactions with human intestinal epithelial cells. Infection and immunity, 80:4089–4098.

Ferrero RL, Lee A. 1988. Motility of campylobacter jejuni in a viscous environment: Comparison with conventional rod-shaped bacteria. J Gen Microbiol, 134:53–59.

Guerry P, Szymanski CM, Prendergast MM, Hickey TE, Ewing CP, Pattarini DL, Moran AP. 2002. Phase variation of campylobacter jejuni 81-176 lipooligosaccharide affects ganglioside mimicry and invasiveness in vitro. Infect Immun, 70:787–793.

Hu L, Tall BD, Curtis SK, Kopecko DJ. 2008. Enhanced microscopic definition of campylobacter jejuni 81-176 adherence to, invasion of, translocation across, and exocytosis from polarized human intestinal caco-2 cells. Infect Immun, 76:5294–5304.

Jan AT. 2017. Outer membrane vesicles (omvs) of gram-negative bacteria: A perspective update. Front Microbiol, 8:1053.

Jang KS, Sweredoski MJ, Graham RLJ, Hess S, Clemons WM. 2014. Comprehensive proteomic profiling of outer membrane vesicles from campylobacter jejuni. J Proteomics, 98:90–98.

Jones MA, Marston KL, Woodall CA, Maskell DJ, Linton D, Karlyshev AV, Dorrell N, Wren BW, Barrow PA. 2004. Adaptation of campylobacter jejuni nctc11168 to high-level colonization of the avian gastrointestinal tract. Infect Immun, 72:3769–3776.

Kelly J, Jarrell H, Millar L, Tessier L, Fiori LM, Lau PC, Allan B, Szymanski CM. 2006. Biosynthesis of the n-linked glycan in campylobacter jejuni and addition onto protein through block transfer. J Bacteriol, 188:2427–2434.

Kidd D, Liu Y, Cravatt BF. 2001. Profiling serine hydrolase activities in complex proteomes. Biochemistry, 40:4005–4015.

Lindmark B, Rompikuntal PK, Vaitkevicius K, Song T, Mizunoe Y, Uhlin BE, Guerry P, Wai SN. 2009. Outer membrane vesicle-mediated release of cytolethal distending toxin (cdt) from campylobacter jejuni. BMC Microbiol, 9:220–220.

Linton D, Allan E, Karlyshev AV, Cronshaw AD, Wren BW. 2002. Identification of n-acetylgalactosamine-containing glycoproteins peb3 and cgpa in campylobacter jejuni. Molecular Microbiology, 43:497–508.

Louwen R, Nieuwenhuis EE, van Marrewijk L, Horst-Kreft D, de Ruiter L, Heikema AP, van Wamel WJ, Wagenaar JA, Endtz HP, Samsom J, van Baarlen P, Akhmanova A, van Belkum A. 2012. Campylobacter jejuni translocation across intestinal epithelial cells is facilitated by ganglioside-like lipooligosaccharide structures. Infect Immun, 80:3307–3318.

Lu Q, Li S, Shao F. 2015. Sweet talk: Protein glycosylation in bacterial interaction with the host. Trends in Microbiology, 23:630–641.

MacCallum A, Haddock G, Everest PH. 2005. Campylobacter jejuni activates mitogen-activated protein kinases in caco-2 cell monolayers and in vitro infected primary human colonic tissue. Microbiology, 151:2765–2772.

Schwechheimer C, Kuehn MJ. 2015. Outer-membrane vesicles from gram-negative bacteria: Biogenesis and functions. Nat Rev Microbiol, 13:605–619.

Szymanski CM, Burr DH, Guerry P. 2002. Campylobacter protein glycosylation affects host cell interactions. Infect Immun, 70:2242–2244.

Szymanski CM, King M, Haardt M, Armstrong GD. 1995. Campylobacter jejuni motility and invasion of caco-2 cells. Infect Immun, 63:4295–4300.

Ternhag A, Torner A, Svensson A, Giesecke J, Ekdahl K. 2005. Mortality following campylobacter infection: A registry-based linkage study. BMC Infect Dis, 5:70.

Tsamandouras N, Chen WLK, Edington CD, Stokes CL, Griffith LG, Cirit M. 2017. Integrated gut and liver microphysiological systems for quantitative in vitro pharmacokinetic studies. AAPS J, 19:1499–1512.

Watson RO, Galan JE. 2008. Campylobacter jejuni survives within epithelial cells by avoiding delivery to lysosomes. PLoS Pathog, 4:e14.

## References

Elmi A, Nasher F, Jagatia H, Gundogdu O, Bajaj-Elliott M, Wren B, Dorrell N. 2016. Campylobacter jejuni outer membrane vesicle-associated proteolytic activity promotes bacterial invasion by mediating cleavage of intestinal epithelial cell e-cadherin and occludin. Cell Microbiol, 18:561–572.

Hall Rl Fau - Miller RJ, Miller Rj Fau - Peatfield AC, Peatfield Ac Fau - Richardson PS, Richardson Ps Fau - Williams I, Williams I Fau - Lampert I, Lampert I. A colorimetric assay for mucous glycoproteins using alcian blue [proceedings].

Liu Y, Patricelli MP, Cravatt BF. 1999. Activity-based protein profiling: The serine hydrolases. Proc Natl Acad Sci U S A, 96:14694–14699.

Valguarnera E, Feldman MF. 2017. Glycoengineered outer membrane vesicles as a platform for vaccine development. Methods Enzymol, 597:285–310.

